# Air mass source determines airborne microbial diversity at the ocean-atmosphere interface of the Great Barrier Reef marine ecosystem

**DOI:** 10.1101/583427

**Authors:** Stephen D.J. Archer, Kevin C Lee, Tancredi Caruso, Katie King-Miaow, Mike Harvey, Danwei Huang, Benjamin J Wainwright, Stephen B Pointing

## Abstract

The atmosphere is the least understood biome on Earth despite its critical role as a microbial transport medium. The influence of surface cover on composition of airborne microbial communities above marine systems is unclear. Here we report evidence for a dynamic microbial presence at the ocean-atmosphere interface of a major marine ecosystem, the Great Barrier Reef, and identify that recent air mass trajectory over an oceanic or continental surface associated with observed shifts in airborne bacterial and fungal diversity. Relative abundance of shared taxa between air and coral microbiomes varied between 2.2-8.8% and included those identified as part of the core coral microbiome. We propose that this variable source of atmospheric inputs may in part contribute to the diverse and transient nature of the coral microbiome.

## Main text

Airborne microbial transport is central to dispersal outcomes [1] and several studies have demonstrated diverse microbial biosignatures are recoverable from the atmosphere. Microbial transport has been shown to occur across inter-continental distances above terrestrial habitats [2–4]. Variation has been recorded seasonally [5, 6], with underlying land use [7], and due to stochastic weather events such as dust storms [8]. Above marine systems the abundance of microorganisms decreases exponentially with distance from land [9], but relatively little is known about potential patterns in biodiversity for airborne microorganisms above the oceans. Here we test the hypothesis that persistent microbial inputs to the ocean-atmosphere interface of the Great Barrier Reef ecosystem vary according to surface cover (i.e. land vs. ocean) during transit of the air-mass.

The Great Barrier Reef is an ideal model system for research on bio-aerosols because incoming air mass during the average residence time for microorganisms in air [10] arises from two distinct sources: a terrestrial continental source in Australia transported across east and northeast dust paths and an oceanic source in the Coral Sea (Fig. 1a). Our study took advantage of a persistent flat-calm sea state during September-October 2016 (www.marineweather.net.au). This minimised interference from microorganisms that are aerosolised by marine spray in heavier sea states at the Great Barrier Reef [11]. We recovered massive bulk-phase air samples at 25m above sea level using a high-volume liquid impinger apparatus (Coriolis μ, Bertin Technologies, France) [12] during a voyage of the *RV Investigator* to circumnavigate the reef (Supplementary Information, Supplementary Methods). We used the National Oceanic and Atmospheric Administration (NOAA) HYSPLIT-WEB model (https://ready.arl.noaa.gov/HYSPLIT.php) to identify back trajectories of air mass during the average residence time for microbial cells in air [10]. Back trajectories for air mass could be delineated clearly into those with recent transit over either continental Australia (continental path) or the Coral Sea (oceanic path) (Fig. 1a; Supplementary Information, Fig. S1). The concurrent concentration of atmospheric radon gas measured in real time was consistently higher from back trajectories originating over continental Australia compared with those that originated from the ocean (Mann-Whitney U Test, P = 0.003). This measurement further validated the binning of air mass into a continental or oceanic origin.

**Figure 1.**
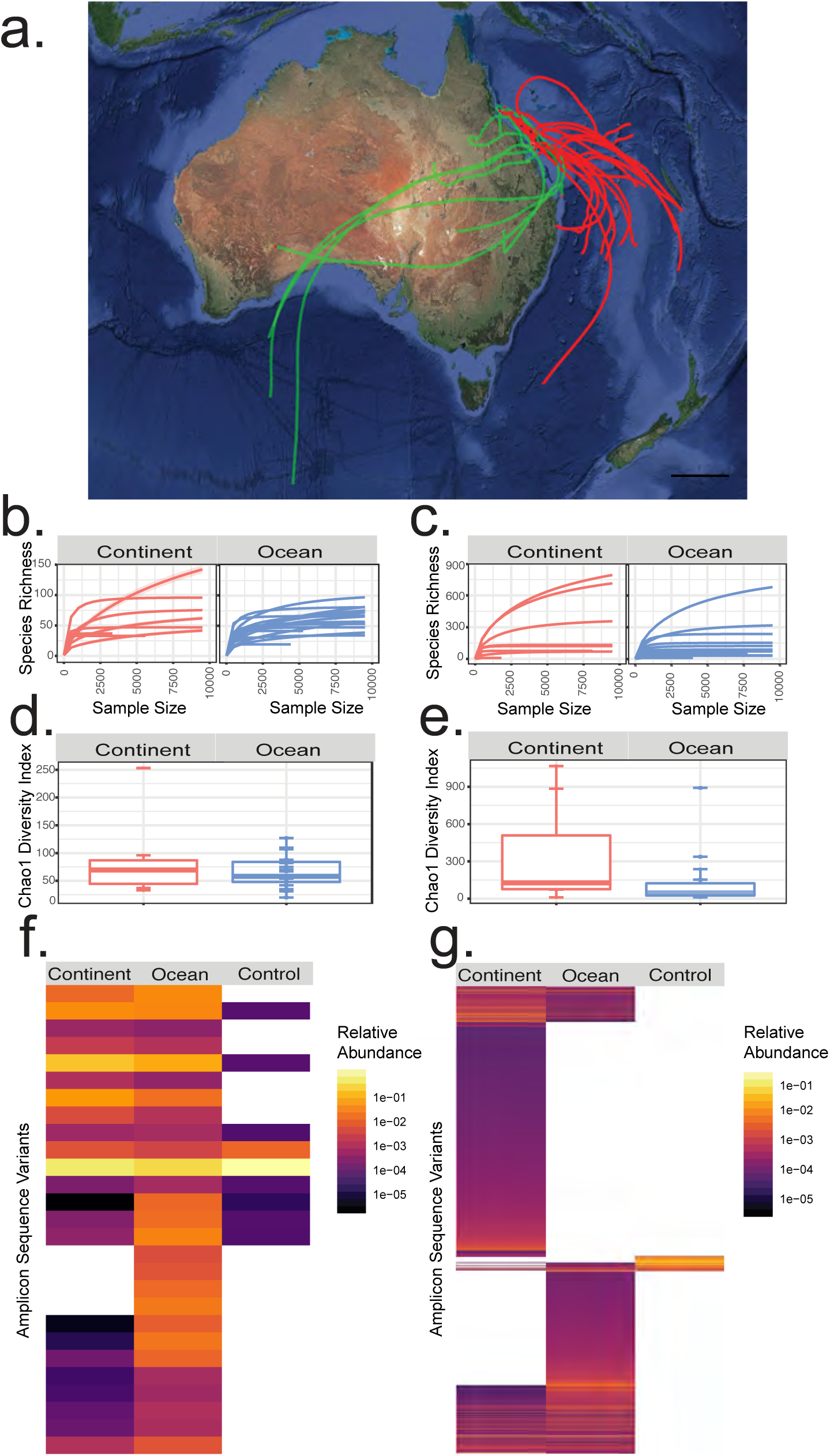
(a) HYSPLIT back trajectory analysis for modelled transit routes (3 d residence time [12]) with colours representing air mass origin over continent (green) or ocean (red) surface, scale bar = 500km; Rarefaction curves for samples by origin for (b) bacteria (Continent n = 8, Ocean n = 19) and (c) fungi (Continent n = 8, Ocean n = 17). Chao1 diversity index by origin for (d) bacteria and (e) fungi. Individual data points are shown as circles, boxplot whiskers represent 1.5 times the interquartile range from the first-third quartiles; (f) Heatmap for bacterial amplicon sequence variants (ASVs) that explain 95% of observed diversity by origin with control samples included (Continent n = 8, Ocean n = 19, control n = 7); (g) Heatmap for fungal ASVs that explain 95% of observed diversity by origin with control samples included (Continent n = 8, Ocean n = 17, control n = 5). All samples were sequenced to near-asymptote. More detailed heatmaps that incorporate the 1,000 most abundant ASVs for bacteria and fungi are shown in Supplementary information, Fig. S3.

Consistent with predictions that microbial biomass is extremely low and unpredictable over the oceans, we achieved recoverable DNA from 27 of 53 bulk-phase air samples (mean recovery 0.23 ng/m^3^, STDEV 0.21 ng/m^3^; Supplementary Information, Table S1) with no significant difference in DNA yield between sample groups (Mann-Whitney U Test, P = 0.37). High-throughput sequencing of the bacterial 16S rRNA gene and fungal Internal Transcribed Spacer (ITS) region were performed as previously described [12] and phylogenetic analysis of amplicon sequence variants (ASVs) were used to estimate diversity [13] (Supplementary Information, Supplementary Methods). Sequencing of control samples revealed very low recovery of putative contaminant microbial signatures. A total of only 17 out of 1403 bacterial and 5 out of 3775 fungal sequences were statistically classified as putative contaminants (Supplementary Information, Supplementary Methods). Most samples were sequenced to near-asymptote (Supplementary Information Fig. 1b, 1c).

Transit above oceanic or continental surfaces was significantly correlated with bacterial and fungal community structures at the ocean-atmosphere interface above the Great Barrier Reef (bacteria: PERMANOVA R^2^ = 0.07233, pseudo F= 1.9494, P = 0.024; manyglm LRT = 1650, P = 0.02; fungi PERMANOVA R^2^ = 0.08413, pseudo F = 2.1125, P = 0.044, manyglm LRT = 6033, P = 0.034). Kendall’s rank correlation tau identified a weak but significant distance relationship (bacteria 0.19 P = <0.001 and fungi 0.10 P = 0.008) (Fig. S5) which suggests air source but also distance can affect community similarities. In terms of overall taxa richness, bacteria displayed similar richness but higher phylogenetic diversity in continental versus ocean-derived samples (Fig. 1d; Supplementary Information Fig. S2), and similarly the fungi displayed markedly greater richness in continent-derived samples (Fig. 1e). Heatmaps of the most abundant ASVs representing 95% of overall diversity illustrate the major taxonomic differences in bacterial (Fig. 1f) and fungal (Fig. 1g) assemblages between ocean and continent-derived air mass. Repeating this analysis with the 1,000 most abundant taxa from each meta-library captured 99.98% of bacterial and 84.24% of fungal diversity in the study and yielded similar results (Supplementary Information Fig. S3). Principal coordinate analysis using Bray-Curtis dissimilarities of bacterial and fungal communities revealed somewhat mixed trends in ordination plots (Supplementary Information Fig. S4), however changes in relative abundance at the phylum level were striking between days when air mass was sourced from continent or ocean (Fig. 2c, 2e).

**Figure 2.**
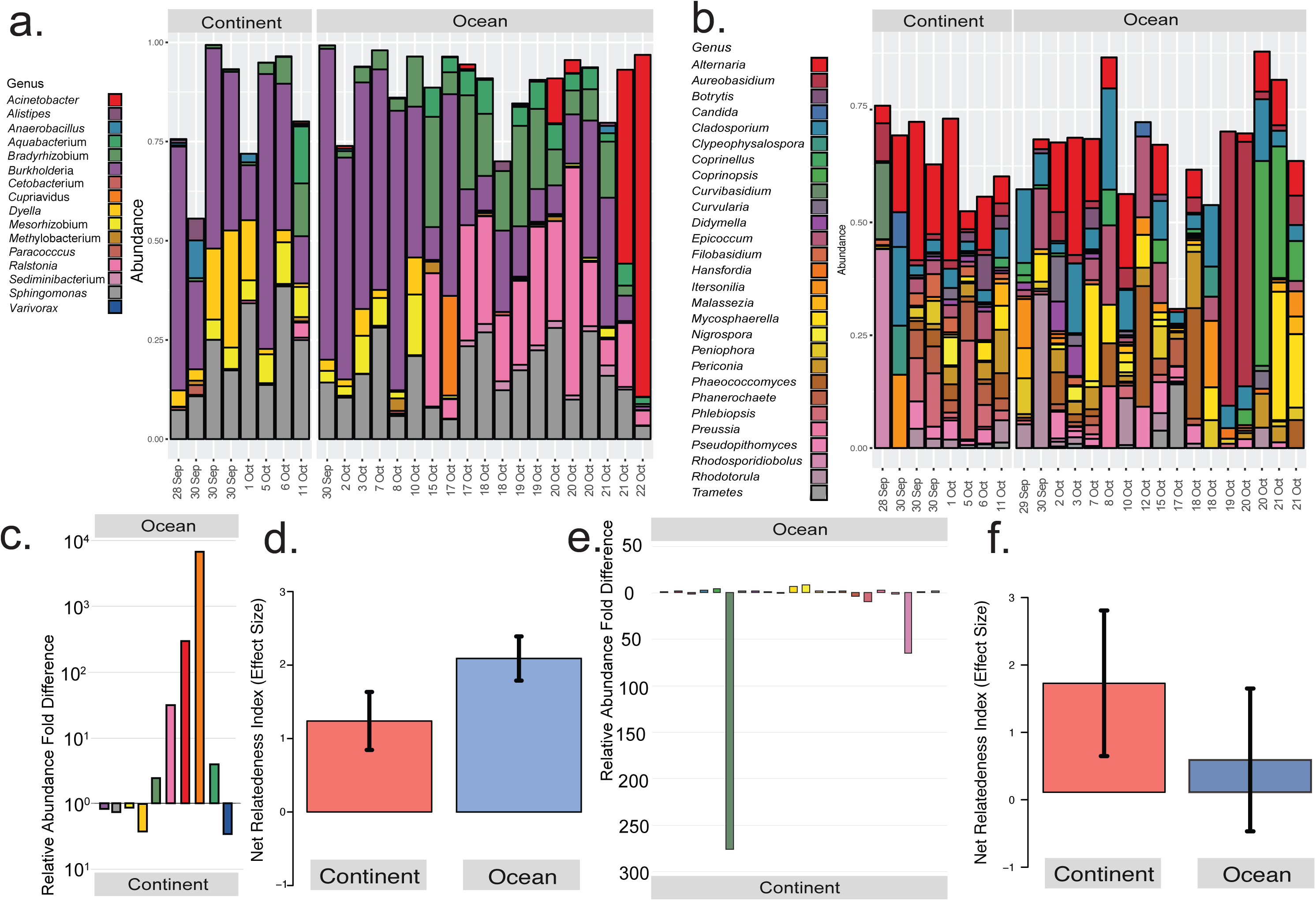
Relative abundance and Genus level taxonomic assignment for ASV of all taxa with >0.5% mean relative abundance for (a) bacteria and (b) fungi; Relative abundance fold differences for taxa from ocean and continent derived air mass, for (c) bacteria and (e) fungi; Net Relatedness Index (NRI) of phylogenetic structure for continental and ocean-derived atmosphere taxa for (d) bacteria(Continent n = 8, Ocean n = 19, control n = 7) and (f) fungal (Continent n = 8, Ocean n = 17, control n = 5) ASVs, error bars indicate standard error of the mean for all samples. Bacterial analysis was based on a phylogenetic tree derived using Maximum Likelihood analysis of 16S rRNA gene ASVs. Fungal analysis utilised a Neighbour Joining tree that was compared with topology of whole genome trees for fungi at high taxonomic ranks, in order to be conservative regarding the phylogenetic information provided by ITS (Supplementary Information, Supplementary Methods).

Overall, the shared ASVs between samples from ocean and continental air masses represented a large proportion of total bacterial (87.9% and 89.6%) and fungal (64.9% and 69.1%) reads. However, the abundances for dominant genera was highly variable between air masses. For example, the three most abundant bacterial genera *Burkholderia*, *Ralstonia* and *Sphingomonas* were present at 28%, 14.4% and 5.5% in oceanic air and 37%, 0.5% and 20% in continental air masses while the three most abundant fungal genera (*Alternaria*, *Cladosporium* and *Mycosphaerella)* were present at 6.4%, 4.5% and 5.3% in oceanic air and 13.5%, 3.5% and 1.4% in continental air masses respectively. A recent study characterizing bacteria aerosolized from a seawater mesocosm concluded approximately 75% of taxa were alpha, beta and gamma Proteobacteria [14], and notably identified *Bradyrhizobium* and *Ralstonia* as highly abundant taxa thus establishing a potential marine source for some of the most abundant airborne taxa in our study. While the core ASVs represented a high proportion of reads, many less abundant ASVs were specific to either oceanic (538 bacteria, 1,335 fungi) or continental (395 bacteria, 1,810 fungi) air masses (Figs. 1f, 1g).

Shifts in relative abundance for bacteria mainly occurred within the phylum Proteobacteria where the dominant genus recovered changed with air mass sampled among *Acinetobacter*, *Alistipes*, *Bradyrhizobium* and *Ralstonia* (Fig. 2a, 2c). All have been recovered as isolates or environmental rRNA gene sequences from both marine and terrestrial sources, thus making any attempt at source tracking challenging. For the Fungi terrestrial air sources supported higher diversity than marine sources and this reflected the abundance of terrestrial fungal habitats. A striking observation was that 56% of recovered genera supported yeast-like taxa, and this provides support for a marine origin since oceanic waters are thought to support elevated abundance of yeasts over filamentous fungi [15]. Shifts in diversity were less pronounced overall for fungi (Fig. 2b, 2d), and were partially obscured by the high diversity relative to bacteria at the genus level. Major shifts in relative abundance occurred for *Aureobasidium, Cladosporium, Coprinopsis*, *Rhodosporidiobolus* and *Rhodotorula*, and all of these genera have known terrestrial and marine records, although it should be noted that terrestrial fungal spores have been recorded in many marine microbial diversity assessments but are unlikely to be active components of an ocean surface water microbiome.

We further interrogated the phylogenetic diversity of continental and oceanic air masses using Net Relatedness Index (NRI) analysis to estimate the level of phylogenetic structuring and putative recruitment from local (i.e. ocean or continental source only) and regional (all sources) pools [12]. The NRI analysis revealed that bacterial assemblages from both oceanic and continental origin displayed positive NRI values with effect sizes indicative of non-random assembly (Fig. 2d, 2f). Communities were thus phylogenetically highly structured, which is possibly due to environmental filtering of traits during transit over the different surface covers. The fungi from continental sources displayed a similar though more variable trend of phylogenetic structuring although in ocean-derived samples this clustering was relatively weak indicating they were more randomly assembled. Overall our analyses support the hypothesis that long-range transport of microbial taxa in air results in differential recruitment and selection during transit over oceanic or continental surfaces [12, 16, 17].

It has been proposed that airborne deposition of microorganisms may be a source of symbionts and pathogenic taxa to coral reefs [18, 19]. In order to establish the potential recruitment of airborne microorganisms to coral reefs we compared our sequence data to that obtained for coral microbiomes. We identified only one study with directly comparable sequence data (Supplementary Information, Supplementary Methods) recovered from *Porites lutea* coral microbiomes in the Andaman Sea and Gulf of Thailand [20]. A total of 6.7% coral reads shared the same ASVs as our airborne bacterial dataset. The air masses from oceanic sources had higher relative abundance of ASVs in common with the coral dataset (8.8%) compared to air with recent continental transit (2.2%) (Supplementary Information, Fig. S6a). Three of the six shared genera (*Bradyrhizobium, Burkholderia* and *Sediminibacterium*) were identified as contributing to the core coral microbiome for the Andaman Sea and Gulf of Thailand [20]. The class Burkholdariales that supports *Burkholdaria* has been identified as a source of ubiquitous coral endosymbionts globally [21]. Relaxing the stringency of this comparison to the Genus level (i.e. ≥97% sequence matches) to account for possible biogeographic variation and ecological variation resulted in higher relative abundance of shared taxa between the datasets. Using this criteria 36% coral bacterial taxa were shared with the atmospheric microbiome, and 77% of oceanic and 79% continental derived atmospheric taxa matched those of the coral microbiome (Supplementary Information, Fig. S6b).

We also compared our bacterial ASV sequences with a recent study of tropical corals from the Singapore Straits, South China Sea [22] which shared significant bacterial 16S rRNA sequence overlaps (Supplementary Information, Supplementary Methods). We identified (with ≥97% sequence identities) 6.5% of coral taxa were shared with the atmospheric microbiome, and 94.4% of ocean derived air and 87.8% of continent derived air taxa were shared with the coral microbiome (Supplementary Information, Fig. S6c). Finally, we broadly screened taxa from our sequence libraries at taxon level (Genus) to known coral-associated taxa including putative symbionts and pathogens identified using different rRNA loci or approaches (e.g. DGGE and Sanger sequencing) (Supplementary Information, Supplementary Methods). Using this approach we estimated the ocean derived air mass supported 8.4% bacterial genera that may include coral associates whilst for continental sources this value was lower at 2.9% (Supplementary Information, Table S2). The most differentially abundant fungal taxon from this comparison (*Ralstonia* sp.) showed greatest sequence similarity to a previously identified coral endosymbiont (Supplementary Information, Table S3) thus indicating any atmosphere-coral microbiome connectivity may be highly variable. For our fungal library there were few coral-associated fungal sequences with which to make comparisons and little phylogenetic information, but at a broad level we identified 8.7% of ocean-derived taxa and 6.2% continent-derived taxa that have recorded coral associates in the same Genus (Supplementary Information, Table S3).

Overall our study has provided a unique insight on the variability of airborne microbial communities above the largest coral reef ecosystem on Earth and yielded clues that atmospheric, oceanic and terrestrial biomes may be inter-connected via the atmospheric microbiome. Our study indicates that the view of coral microbiomes as harbouring few long term residents and instead comprising largely a “diverse transient community that is responsive to surrounding environment”[23] may be explained in part by the variability in airborne microbial diversity above reefs that may act as recruitment reservoirs. Improved understanding of cross-biome microbial biocomplexity and interaction will require coordinated and directly comparable research effort from atmospheric, marine and terrestrial microbiologists.

## Data availability

All sequence data generated by this study has been submitted to the EMBL European Nucleotide Archive (ENL) under BioProject PRJEB31630 with sample accession numbers ERS3215240 to ERS3215312.

## Author contribution statement

S.D.J.A, K.C.L, T.C., M.H. and S.B.P. conceived the study; S.D.J.A. conducted ship-board fieldwork; S.D.J.A and K.C.L. performed laboratory experiments; S.D.J.A., K.C.L., T.C., K.K-M., B.J.W. and S.B.P. performed data analysis; S.D.J.A, M.H., D.H., B.J.W. and S.B.P. analysed and interpreted the findings; S.B.P. wrote the manuscript.

## Acknowledgements

Collaboration in this work was supported through ARC Discovery Project (grant number DP150101649) and NIWA’s Research Programme in Ocean-Climate Interactions (grant number 2017/18 SCI). Financial support for laboratory research was provided by the Yale-NUS College Start-up Fund. Author SDJA was supported by a Postdoctoral Fellowship from Auckland University of Technology (AUT). We thank ARC Discovery Project Principal Investigators Prof. Zoran Ristovski and Luke Cravigan (Queensland University of Technology), the officers and crew of the RV Investigator (IN2016_V05), and Tony Bromley and Sally Gray (NIWA) for their outstanding logistical support and Timothy Lawrence (AUT) for his DNA sequencing support. Radon data was provided by Scott Chambers and Alastair Williams (Australian Nuclear Science and Technology Organisation).

