## Supplementary Information for "Air mass source determines airborne microbial diversity at the ocean-atmosphere interface of the Great Barrier Reef marine ecosystem"

### Supplementary Methods

#### Sample Recovery

Our study took advantage of a persistent flat-calm sea state between 30th September to 22nd October 2016 (www.marineweather.net.au) that minimised interference from microorganisms that are aerosolised by marine spray in heavier sea states at the Great Barrier Reef. The voyage of the *RV Investigator* circumnavigated the reef. We sampled air mass daily from the deck of the research vessel approximately 25m above the sea surface. Massive bulk-phase air volumes (18-108 $m^3$ per sample) were collected for each discrete sampling station. Overall, we retrieved 53 independent bulk-phase air samples (Supplementary Information, Table S1).

We employed a high-volume liquid impinger apparatus (Coriolis μ, Bertin Technologies, France) and collected aerosol particulate fractions directly into phosphate-buffered saline (PBS) using collection cones supplied by the manufacturer. Evaporation was compensated for using a peristaltic pump set to between 0.5 and 1.3 mL per minute with PBS. Random collection cones were also assembled into the machine but not activated, and these were used as the negative controls. All collection cones were soaked in 1.5 % sodium hypochlorite (NaClO) then washed with 70 % ethanol and three washes of Milli-Q $H_20$ before being filled with filtered PBS. All sampling equipment was disassembled between locations and cleaned with NaClO, ethanol and Milli-Q $H_20$. Samples were processed immediately on board the vessel.

Back-trajectories of air mass arriving at each sampling interval were generated using the National Oceanic and Atmospheric Administration (NOAA) HYSPLIT-WEB model (https://ready.arl.noaa.gov/HYSPLIT.php) (Fig. S1). Calculations were made with a 3 day back trajectory because this is estimated as the average residence time for microbial cells in air [1] (Fig. S1). We acknowledge that large variation is inherent in models of atmospheric transport and uncertainty increases with duration of the back trajectory estimate. With our model the estimated trajectories greater than two weeks resulted in diffuse origins for all samples from above the Southern Ocean. The HYSPLIT back trajectories were calculated using the GDAS database and the model vertical velocity option. The continental or oceanic origin of air mass was further verified by measuring the concurrent concentration of atmospheric radon gas on board the research vessel in real time by the Australian Nuclear Science and Technology Organisation (ANSTO) (Supplementary Information, Table S1) [2], with the hypothesis that radon levels are typically higher from back trajectories originating over continental Australia compared with those that originate from the ocean. Radon is emitted only from unsaturated, unfrozen terrestrial surfaces and has a half-life of 3.8 days, making it a powerful tracer of continental air transport and mixing.

**Environmental DNA sequencing and bioinformatics**

Airborne microbial samples suspended in PBS were filtered onto a 25 mm 0.2 μm polycarbonate filter and stored frozen until processed. Total DNA was directly extracted using a CTAB protocol [3]. DNA yield for these ultra-low biomass samples was quantified using the Qubit 2.0 Fluorometer (Invitrogen) (Supplementary Information, Table S1). Samples were then stored at -20 °C until processed. We used DNA yield as an indirect estimate for biomass.

Illumina MiSeq libraries were prepared as per manufacturer's protocol (Metagenomic Sequencing Library Preparation Part # 15044223 Rev. B; Illumina, San Diego, CA, USA) and as previously described with PhiX positive controls [4]. We targeted bacteria and fungi since these domains are the most abundant microorganisms in aerosols [5]. PCR was conducted with primer sets targeting the V3-V4 regions of bacterial and archaeal 16S rRNA gene: PCR1 forward (5' TCGTCGGCAG CGTCAGATGT GTATAAGAGA CAGCCTACGG GNGGCWGCAG 3') and PCR1 reverse (5' GTCTCGTGGG CTCGGAGATG TGTATAAGAG ACAGGACTAC HVGGGTATCT AATCC 3') and the internal transcribed spacer region of fungal 18S and 5.8S rRNA genes: ITS1 forward (5' CTTGGTCATTTAGAGGAAGTAA 3') and ITS2 reverse (5' GCTGCGTTCTTCATCGATGC 3'). These primers for Bacteria and Fungi are widely accepted to capture the broadest estimates of diversity [6–8]. Total sequence library sizes were 12,027,554 for bacteria and 8,570,900 for fungi before filtering. Total counts in the processed dataset were 4,727,038 bacterial and 2,969,686 fungal sequences. A total of 1386 bacterial and 3775 fungal taxa were identified from these.

Sequencing data for bacterial 16S rRNA gene and fungal ITS1 amplicons was processed based on the DADA2 v1.8 [9] pipeline. Primers sequences were removed using cutadapt [10] to remove forward (CCTACGGGNGGCWGCAG) and reverse (GACTACHVGGGTATCTAATCC) for 16S rRNA gene, and forward (CTTGGTCATTTAGAGGAAGTAA) and reverse (CTTGGTCATTTAGAGGAAGTAA) for fungal ITS1 region. Bacterial reads were uniformly trimmed to 280 bp (forward) and 250 bp (reverse) and then filtered by removing reads exceeding maximum expected error of 2 for forward reads and 5 for reverse reads or reads containing ambiguity N symbol. Fungal reads were untrimmed (due length hyper-variability of the region) and reads exceeding maximum expected error of 5 for forward reads and 8 for reverse reads, or those with ambiguity N symbols were removed. The reads were used to train the error model and then dereplicated to acquire unique sequences, which were used to infer sequence variants with the trained error model. The forward and reverse reads were merged and chimeric sequences were removed. For bacteria we used amplicon sequence variants (ASVs) to assign taxa since this has been shown as the most robust method currently available for bacterial 16S rRNA gene-defined taxa identification [11]. ASVs were given taxonomic assignment using DADA2 with SILVA nr v132 database [12] for bacterial reads to provide species level assignment based on exact match between ASVs and known reference sequences. . Fungal reads were taxonomically classified using UNITE v7.2 database [13]. All sequence data generated by this study has been submitted to the EMBL European Nucleotide Archive (ENL) under BioProject PRJEB31630 with sample accession numbers ERS3215240 to ERS3215312.

The resulting taxa were then processed as previously described [14] [15]. The R packages phyloseq [16], DESeq2 [17] and ggplot2 [18] were used for downstream analysis and visualisation including ordination and alpha/beta diversity calculations. For heatmap visualisations the 1,000 most abundant ASVs in each dataset were selected and these captured 99.99 % bacterial and 91.74 % fungal reads in the libraries. All other analyses used the entire sequence library data. Our heatmap analysis therefore had high confidence since the unsampled 'tail' comprised only extremely rare sequence variants at very low/singleton abundance.

Sequencing of control samples revealed very low recovery of putative contaminant microbial signatures, a total of only 17 out of 1403 bacterial and 5 out of 3775 fungal sequences were statistically classified as putative contaminants with *decontam* 1.2.0 [19].

**Meta-analysis of atmospheric microbiome with coral microbiomes**

A meta-analysis of our data with publicly available sequence data from coral-associated microbiomes was conducted. We first searched for relevant recent published studies and constrained our search as follows: The first 20 matches by relevance from the search parameters ["16S" "coral"] and ["16S" "reef"] from 2015 to 2019 with Google

Scholar were employed to determine if there was any publicly available next-generation sequencing data that encompassed the compatible 16S rRNA gene region. Based on these criteria, we identified a single publication [20] which allowed for direct ASV analysis. Another dataset with limited overlap was also assessed based on similarity searches of representative sequences to determine shared taxa [21]. Some excellent studies on Great Barrier Reef coral microbiomes could not be included because they have employed the V1-V3 region of the 16S rRNA gene for bacteria and this does not overlap with the V3-V4 region used in our study. We also searched nucleotide databases (NCBI and EMBL) for unpublished but publicly available coral microbiome sequences using the same search criteria. This search revealed three entries in the top 50 matches by relevance but our initial quality checking revealed these were unsuitable for further inclusion in our analysis.

PacBio SMRT sequencing of 16S rRNA gene amplicons from corals of Gulf of Thailand and Andaman Sea was conducted by Pootakham et al., (2017)[20]. The raw reads were retrieved from NCBI Short Read Archive (accession SRR5149735) via NCBI SRA toolkit (https://github.com/ncbi/sra-tools/wiki). As the reads from this study extend beyond those of our study (V3-V4), we were able to process the reads in a manner similar to our study and trim the reads based on the primer regions of our study (341F/785R). This resulted in reads targeting homologous regions for comparison between the two studies. The trimmed reads followed a similar processing pipeline as our study (dada2 for ASV-calling and taxonomic classification, see below), which enabled direct and precise comparison of ASV sequences.

The Wainwright et al., (2019) study [21] conducted paired-end Illumina sequencing of 16S rRNA gene amplicons of the coral *Pocillopora acuta* around Singapore. The coral study utilised comparable data processing pipeline (specifically dada2 to generate ASV) but used a different set of primers to amplify 16S rRNA gene (V4 vs our V3-V4 region). However, a significant portion of the partial 16S rRNA gene amplicon still overlapped (~250 bp). The ASVs from our study were searched against reference ASVs(https://github.com/gzahn/Pocillopora_Bacteria) from Wainwright et al. (2019) using the usearch v10.0.240_i86linux32 global function [22]. Both strand orientations were searched (-strand both) and a minimum identity threshold (-id) of 0.97 was set. The ASVs were sorted with descending number in relation to mean relative abundance (i.e. ASV1 has highest mean relative abundance).

We also conducted a more relaxed analysis where the 100 most abundant bacterial and fungal taxa at genus level (Table S1-3) were compared to published studies that used different approaches for coral microbiome diversity estimation [23–35]. Microorganisms unassigned at the Genus taxonomic level were excluded from further analysis. A weighted average of the relative abundance was calculated for each ASV identified in this study, grouping samples with an ocean air mass source, and a continental air mass source. Near full length 16S rRNA gene sequences of coral associated bacteria were searched against our ASV sequences using usearch_local function in usearch v10.0.240 to identify highly similar matches.

For the fungi no comparable sequence data for coral-associated fungi is available. We therefore performed an approximate comparison at the Genus level with reference to published descriptions of coral-associated fungi [23, 24, 28, 29]. There are no bacterial or fungal taxa that have yet been unequivocally been identified as either obligate symbionts [27] or pathogens [36] of corals.

**Statistical Treatments**

Community phylogenetic metrics including were calculated using the R [37] packages picante [38], ape [39], phylobase [40], adephylo [41], and phytools [42]. We tested partitioning of Bray-Curtis dissimilarity matrices by air mass origins (continent or ocean) through PERMANOVA, using the *adonis* function from R package vegan 2.5-5 [43] with 999 unrestricted permutations. Phylogenetic trees for community phylogenetic structure analysis were constructed for all OTUs with FastTree v2.1.9 on multiple alignment of sequences produced by MUSCLE v3.8.31. For Bacteria an approximately Maximum-Likelihood approach was used whilst for Fungi an alignment-free distance approach with Neighbour-Joining method was employed in order to generate a ITS-based phylogenetic tree for community metrics [44]. The distance approach was used for Fungi as the hypervariability of ITS1 loci hinders multiple sequence alignment required for most phylogenetic analyses. To validate this approach, we compared our tree topology with the most recent whole genome phylogenies for the Fungi [45, 46]. This approach was robust at higher taxonomic levels (there is no consensus for fungal phylogenies using multiple loci or whole genomes below Order rank) and has been used successfully with other eukaryotic taxa as a workflow for Net Relatedness analysis [47]. Although we acknowledge limitations to this approach, the advantages of using ITS loci for taxonomic identification vastly outweighed its shortcomings, and we also triangulated data from this test with additional analytical approaches, so overall our inclusion of this test is justified and interpreted conservatively. Mean phylogenetic distance (MPD) [48] was calculated to measure phylogenetic distance between ASVs and OTUs in each sample. A null model algorithm based on independent swap (999 randomisations) was used to test the extent of phylogenetically clustering (positive values for the NRI effect size) or over-dispersion (negative values for the NRI effect size) [49].

**Methods References**

18. Wickham H. ggplot2. 2009. Springer New York, New York, NY.

35. Nelson CE, Alldredge AL, Mccliment EA, Amaral-zettler LA, Carlson CA. Depleted dissolved organic carbon and distinct bacterial communities in the water column

of a rapid-flushing coral reef ecosystem. *ISME J* 2011; **5**: 1374–1387.

36. Rosenberg E, Koren O, Reshef L, Efrony R, Zilber-Rosenberg I. The role of microorganisms in coral health, disease and evolution. *Nat Rev Microbiol* 2007; **5**: 355–362.

37. Swenson NG. Functional and Phylogenetic Ecology in R. 2014. Springer New York, New York, NY.

**Figure S1.** HYSPLIT back trajectory analysis for modelled transit routes with sampling date indicated (full 3 day back trajectories are shown in Fig. 1).

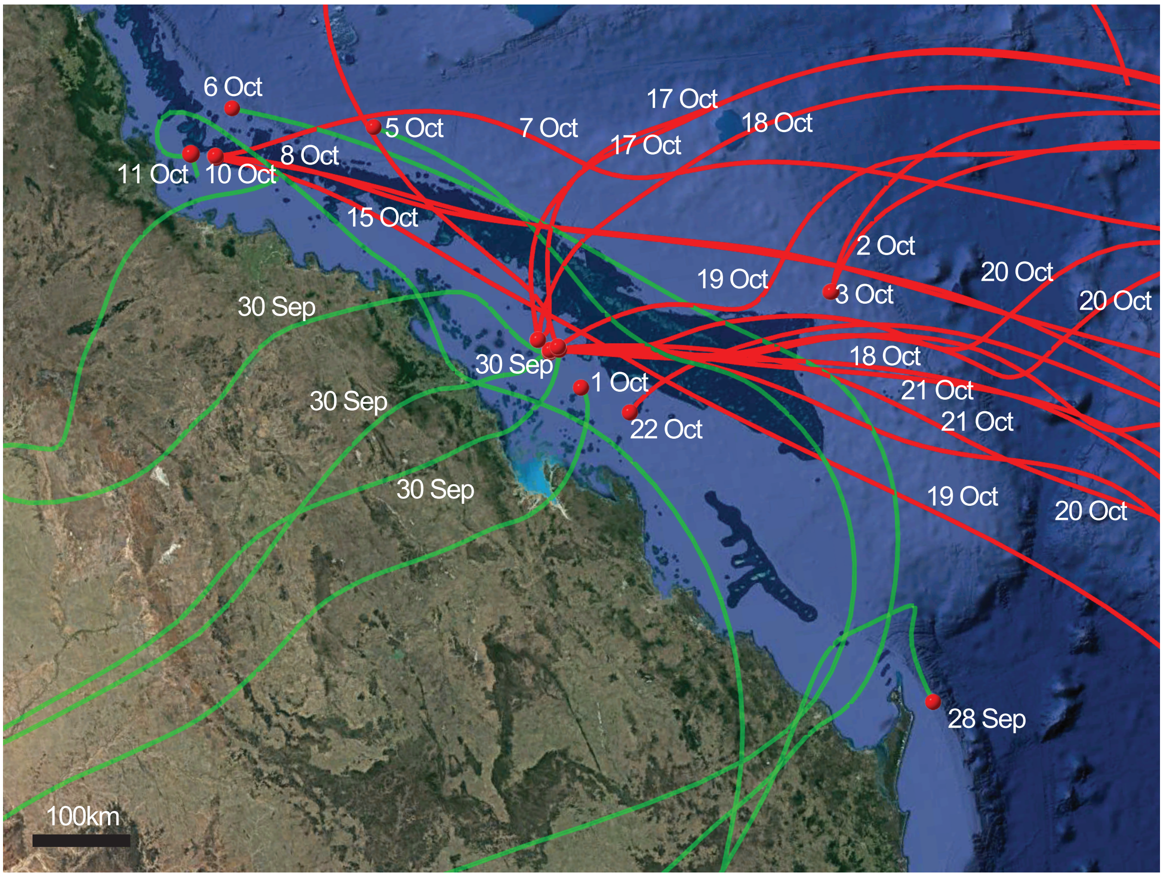

**Table S1.** Sampling meta-data.

| SAMPLE NAME | Long. (decimal degrees) | Lat. (decimal degrees) | Radon (mBq/m$^3$) | Air Source | Date Start | Time Start | Duration (hrs) | Volume (M$^3$) | Total DNA ng/M$^3$ | Passed Sequencing (Y/N) |
|---|---|---|---|---|---|---|---|---|---|---|
| **GBR-01-SA** | 24.8973 | 153.5982 | 1086.1301 | continent | 28.09.16 | 1500 | 4 | 72 | below detection | Y |
| **GBR-02-SA** | na | na | na | na | 29.09.16 | 1000 | 4 | 72 | 0.075 | N |
| **GBR-03-1-SA** | -20.9937 | 150.1999 | error | ocean | 30.09.16 | 0800 | 0.5 | 9 | 0.244 | Y |
| **GBR-04-SA** | na | na | na | na | 30.09.16 | 1100 | 4 | 72 | 0.722 | N |
| **GBR-05-1** | -20.9984 | 150.2017 | error | continent | 30.09.16 | 1500 | 4 | 72 | below detection | Y |
| **GBR-05-2** | -20.9947 | 150.2022 | error | continent | 30.09.16 | 1900 | 4 | 72 | 0.037 | Y |
| **GBR-05-3** | -21.0264 | 150.2081 | error | continent | 30.09.16 | 2300 | 4 | 72 | below detection | Y |
| **-VE CONTROL 1** | na | na | na | control | | | | 0 | below detection | Control |
| **GBR-05-4** | 21.4199 | 150.3747 | 1549.0931 | continent | 01.10.16 | 0300 | 4 | 72 | below detection | Y |
| **GBR-05-5** | na | na | na | na | 01.10.16 | 0700 | 4 | 72 | 1.467 | N |
| **GBR-05-6** | na | na | na | na | 01.10.16 | 1100 | 4 | 72 | 0.111 | N |
| **GBR-06-1** | na | na | na | na | 2.10.16 | 0730 | 6 | 108 | below detection | N |
| **GBR-06-2** | na | na | na | na | 2.10.16 | 1330 | 6 | 108 | below detection | N |
| **GBR-06-3** | -20.8275 | 153.0668 | 499.6199 | ocean | 2.10.16 | 1930 | 6 | 108 | below detection | Y |
| **GBR-06-4** | -20.8251 | 153.0786 | 552.5687 | ocean | 3.10.16 | 0130 | 2 | 36 | 0.199 | Y |
| **-VE CONTROL 2** | na | na | na | control | | | | 0 | below detection | Control |
| **GBR-07-SA** | -18.5789 | 148.6524 | 904.2033 | continent | 5.10.16 | 0900 | 6 | 108 | 0.118 | Y |
| **GBR-08-SA** | -18.1407 | 147.2207 | 434.4520 | continent | 6.10.16 | 0900 | 6 | 108 | 0.081 | Y |
| **GBR-09-SA** | -18.5786 | 146.9731 | 389.6492 | ocean | 7.10.16 | 0900 | 6 | 108 | below detection | Y |
| **GBR-09-2** | na | na | na | na | 7.10.16 | 1500 | 6 | 108 | below detection | N |
| **GBR-10-SA** | 18.5847 | 146.9712 | 210.4377 | ocean | 8.10.16 | 0900 | 6 | 108 | below detection | Y |
| **GBR-11-1** | na | na | na | na | 9.10.16 | 0900 | 6 | 108 | below detection | N |
| **GBR-11-2** | na | na | na | na | 9.10.16 | 1500 | 6 | 108 | below detection | N |
| **-VE CONTROL 3** | na | na | na | control | | | | 0 | below detection | Control |
| **GBR-11-3** | na | na | na | na | 9.10.16 | 2100 | 6 | 108 | 0.070 | N |
| **GBR-11-4** | 18.5822 | 146.9790 | 205.0071 | ocean | 10.10.16 | 0300 | 6 | 108 | 0.046 | Y |
| **GBR-11-5** | na | na | na | na | 10.10.16 | 0900 | 6 | 108 | 0.039 | N |
| **GBR-12-SA** | 18.5164 | 146.7233 | 811.8823 | continent | 11.10.16 | 0900 | 6 | 108 | 0.039 | Y |
| **GBR-13-SA** | na | na | na | na | 12.10.16 | 0900 | 6 | 108 | below detection | N |
| **GBR-13-2** | na | na | na | na | 12.10.16 | 1500 | 6 | 108 | below detection | N |
| **GBR-13-3** | na | na | na | na | 13.10.16 | 0230 | 3.5 | 63 | below detection | N |
| **-VE CONTROL 4** | na | na | na | control | | | | 0 | below detection | Control |
| **GBR-14-SA** | 18.6035 | 146.9800 | 279.6785 | ocean | 15.10.16 | 1300 | 4 | 72 | 0.028 | Y |
| **GBR-15-1** | 20.9020 | 150.0082 | 270.1749 | na | 17.10.16 | 1400 | 1 | 18 | below detection | N |
| **GBR-15-2** | na | na | na | na | 17.10.16 | 1800 | 1 | 18 | below detection | N |
| **GBR-15-3** | 20.9019 | 150.0029 | 268.8172 | na | 17.10.16 | 2200 | 1 | 18 | below detection | N |
| **GBR-15-4** | na | na | na | na | 18.10.16 | 0200 | 1 | 18 | below detection | N |
| **GBR-15-5** | 20.8992 | 150.0069 | 243.0216 | na | 18.10.16 | 0600 | 1 | 18 | below detection | N |

| | | | | | | | | | | |
|---|---|---|---|---|---|---|---|---|---|---|
| **GBR-15-6** | na | na | na | na | 18.10.16 | 1000 | 1 | 18 | below detection | N |
| **-VE CONTROL 5** | na | na | na | control | | | | 0 | below detection | Control |
| **GBR-15-7** | na | na | na | na | 18.10.16 | 1400 | 1 | 18 | below detection | N |
| **GBR-15-8** | 21.0396 | 150.1081 | error | na | 18.10.16 | 1800 | 1 | 18 | below detection | N |
| **GBR-15-9** | na | na | na | na | 18.10.16 | 2200 | 1 | 18 | below detection | N |
| **GBR-15-10** | na | na | na | na | 19.10.16 | 0200 | 1 | 18 | below detection | N |
| **GBR-15-11** | 21.0292 | 150.1081 | 205.0071 | ocean | 19.10.16 | 0600 | 1 | 18 | below detection | Y |
| **GBR-15-12** | 21.0221 | 150.1135 | 272.8902 | ocean | 19.10.16 | 1000 | 1 | 18 | below detection | Y |
| **GBR-16-1** | 21.0151 | 150.1145 | 380.1455 | ocean | 20.10.16 | 0600 | 1 | 18 | below detection | Y |
| **-VE CONTROL 6** | na | na | na | control | | | | 0 | below detection | Control |
| **GBR-16-2** | na | na | na | na | 20.10.16 | 1000 | 1 | 18 | below detection | N |
| **GBR-16-3** | 21.0187 | 150.1148 | 716.8459 | ocean | 20.10.16 | 1400 | 1 | 18 | below detection | Y |
| **GBR-16-4** | 21.0183 | 150.1139 | 541.7074 | ocean | 20.10.16 | 1800 | 1 | 18 | below detection | Y |
| **GBR-16-5** | na | na | na | na | 20.10.16 | 2200 | 1 | 18 | below detection | N |
| **GBR-16-6** | na | na | na | na | 21.10.16 | 0200 | 1 | 18 | 0.236 | N |
| **GBR-16-7** | 21.0173 | 150.1127 | 427.6637 | ocean | 21.10.16 | 0600 | 1 | 18 | below detection | Y |
| **GBR-16-8** | 21.0168 | 150.1131 | 389.6492 | ocean | 21.10.16 | 1000 | 1 | 18 | below detection | Y |
| **-VE CONTROL 7** | na | na | na | control | | | | 0 | below detection | Control |
| **GBR-16-9** | na | na | na | na | 21.10.16 | 1400 | 1 | 18 | 0.147 | N |
| **GBR-16-10** | na | na | na | na | 21.10.16 | 1800 | 1 | 18 | below detection | N |
| **GBR-16-11** | na | na | na | na | 21.10.16 | 2200 | 1 | 18 | below detection | N |
| **GBR-16-12** | 21.7245 | 150.8378 | 271.5325 | ocean | 22.10.16 | 0200 | 1 | 18 | below detection | Y |

**Figure S2.** Faith's Phylogenetic Diversity (PD) for bacterial ASV originating from continental or oceanic derived air masses, with Chao1 diversity estimates shown for comparison. Phylogenetic diversity could not be performed with fungal ITS1 sequences due to the hypervariability of the region prohibiting the construction of a reliable phylogenetic tree.

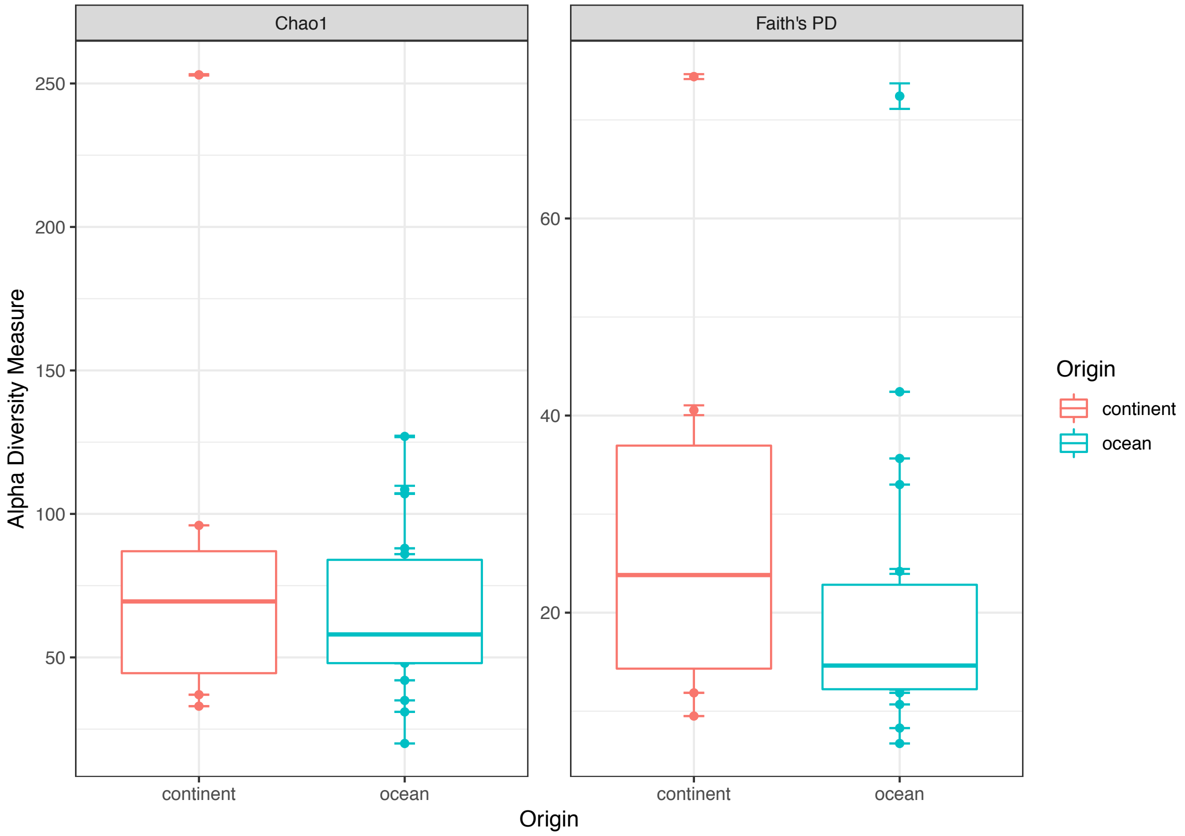

**Figure S3.** (a) Heatmap for 1,000 most abundant bacterial amplicon sequence variants (ASVs) by origin with control samples included (Continent n = 8, Ocean n = 19, control n = 7); (b) Heatmap for 1,000 most abundant fungal ASVs by origin with control samples included (Continent n = 8, Ocean n = 17, control n = 5). All samples were sequenced to near-asymptote. This analysis captured 99.98% of bacterial and 84.24% of fungal overall ASV-defined diversity.

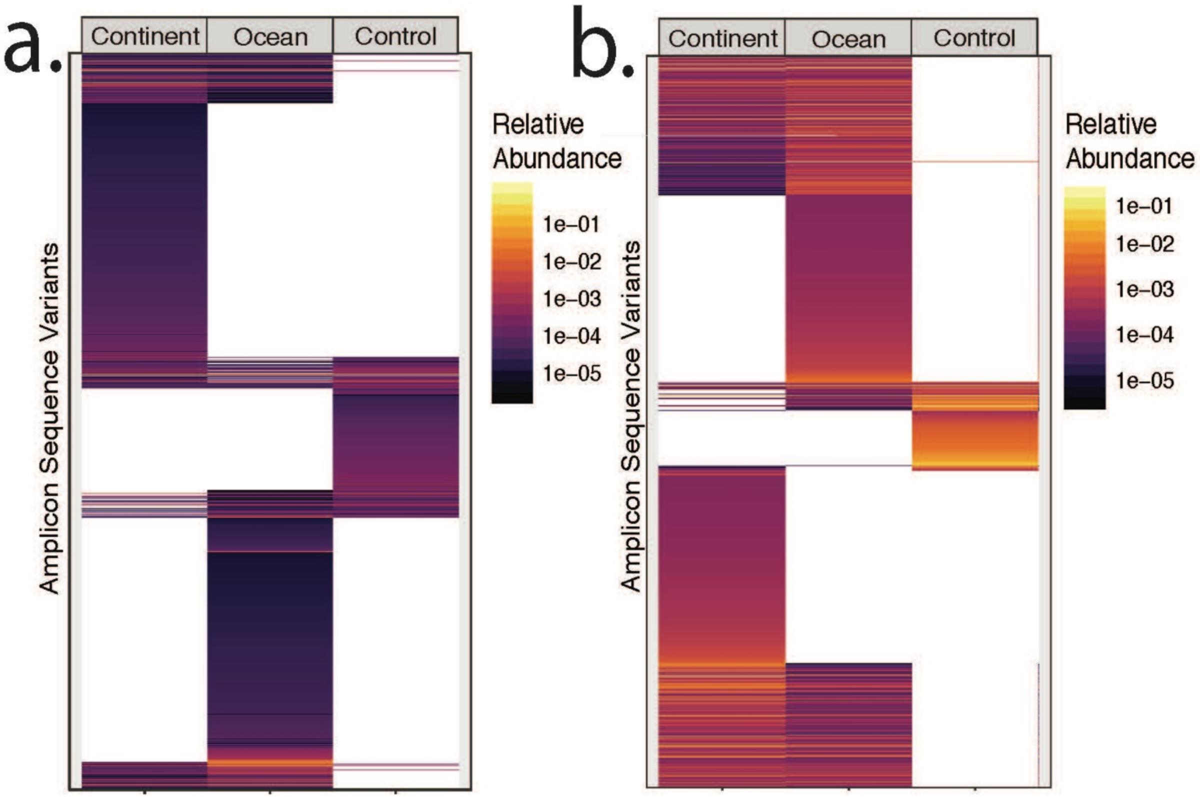

**Figure S4**. PCoA of Bray Curtis Dissimilarities for (a) bacteria and (b) fungi.

a.

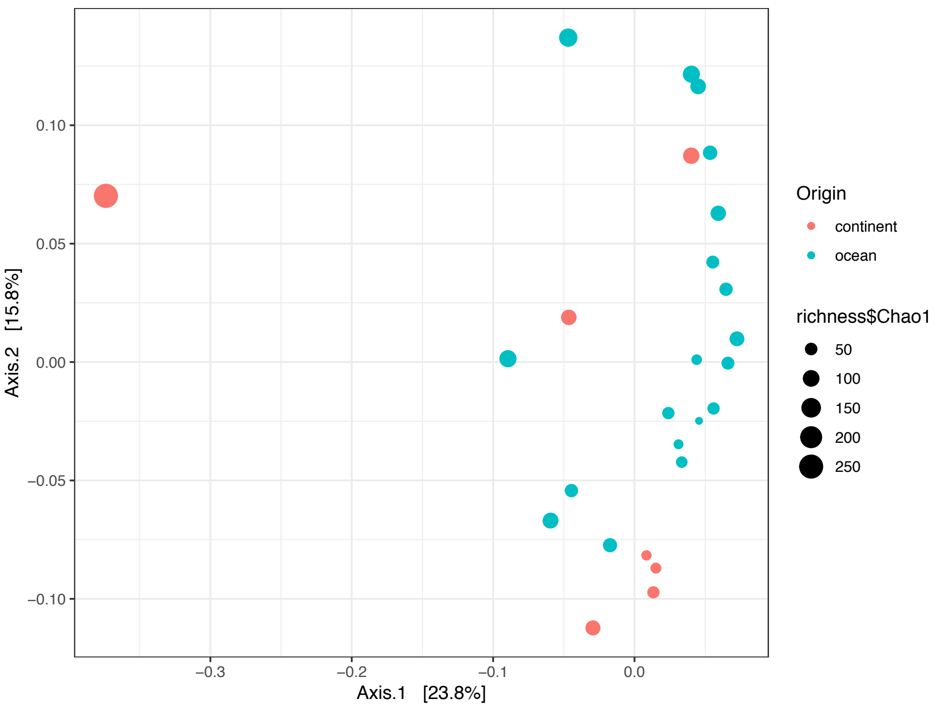


b.

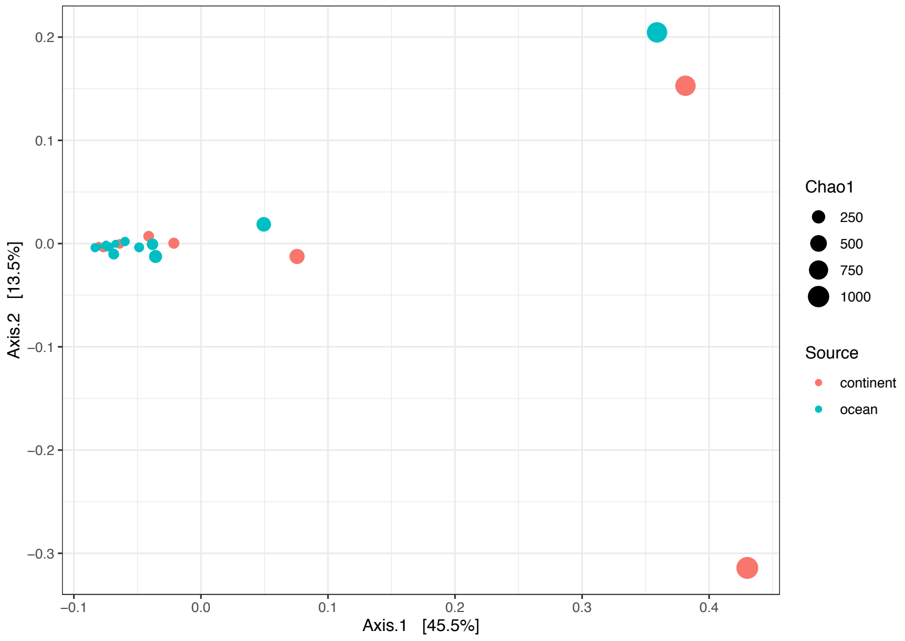

**Figure S5.** Linear distance decay using Kendall's rank correlation tau for (a) bacteria and (b) fungi.

a)

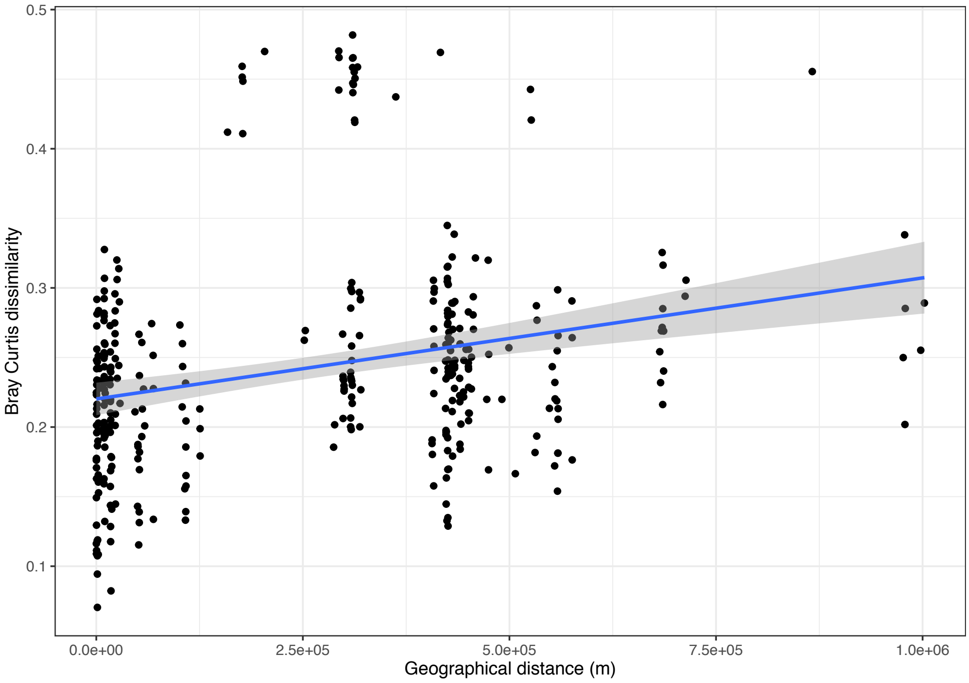


b)

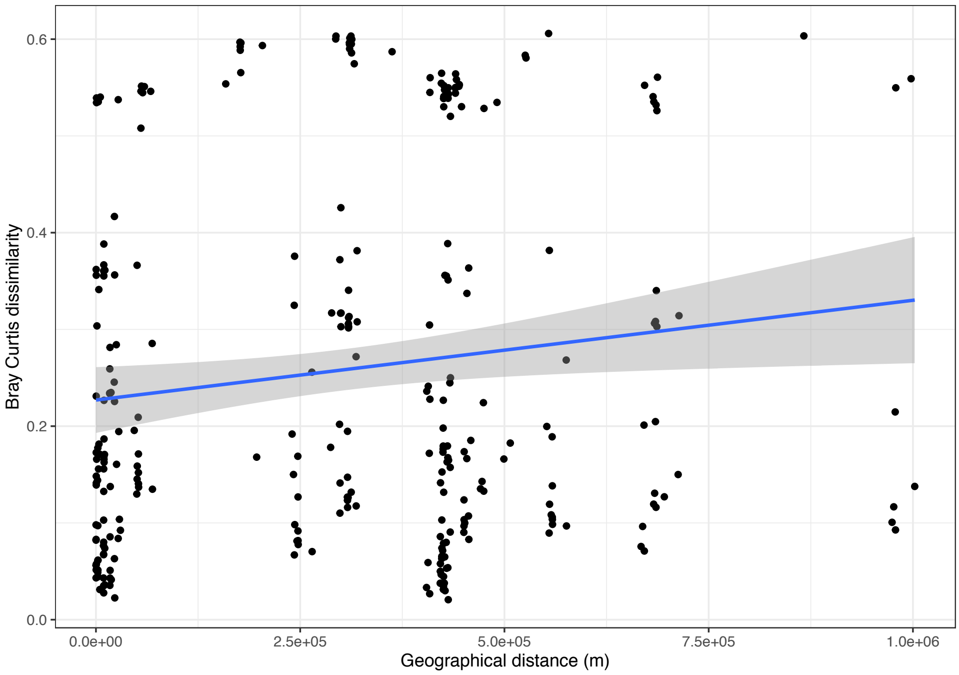

**Figure S6.** Relative abundance of bacterial taxa between air and coral sequence libraries for all taxa with >0.05% mean relative abundance. a) direct ASV matches between our study by source and the Pootakham et al coral microbiome [20], b) direct ASV matches that are classified at genus level between our study and the Pootakham et al coral microbiome [20] c) shared ASVs (by ≥97% sequence identity) between our study and the Wainwright et al coral microbiome [21].

.

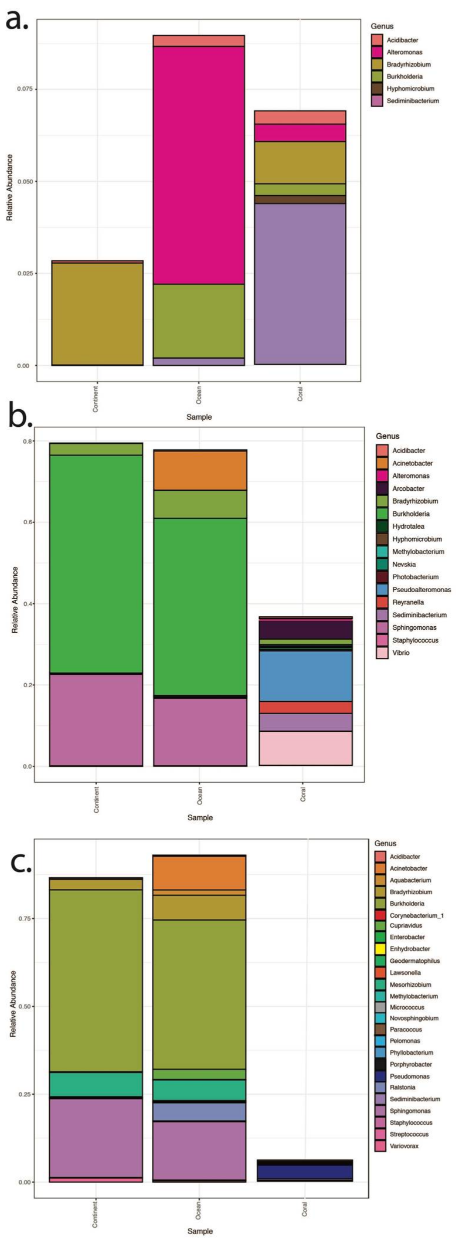

**Table S2.** Relative abundances of the 100 most abundant bacterial ASV's from genera identified in the literature as components of coral associated microbiomes**.**

| **Publication** | **Sample location** | **Microbiome type** | **Total number of matching genera** | **Matching genera** | **Abundance continent from matching genera** | **Abundance ocean matching genera** |
|---|---|---|---|---|---|---|
| Angly et al.<br>(2016)<br>doi: 10.7717/peerj.1511 | Great Barrier Reef | Water associated from reef surrounds | 12 | *Sphingomonas*<br>*Acinetobacter*<br>*Vibrio*<br>*Bacillus*<br>*Novosphingobium*<br>*Clostridium*<br>*Flavobacterium*<br>*Cellvibrio*<br>*Pseudoalteromonas*<br>*Pseudomonas*<br>*Lysobacter*<br>*Cupriavidus* | 0.2255 | 0.2877 |
| Bourne and Munn<br>(2005)<br>doi:10.1111/j.1462-2920.2005.00793.x | Great Barrier Reef | Coral associated | 2 | *Vibrio*<br>*Shewanella* | ND | ND |
| Cardinas et al.<br>(2011)<br>doi: 10.1038/ismej.2011.123 | Caribbean - Colombia | Coral associated | 9 | *Vibrio*<br>*Pantoea*<br>*Pseudomonas*<br>*Stenotrophomonas*<br>*Paracoccus*<br>*Bacillus*<br>*Microbacterium*<br>*Brevibacterium*<br>*Micrococcus* | ND | ND |
| Sato et al.<br>(2009)<br>doi:10.1038/ismej.2009.103 | Great barrier reef | Coral associated and pathogens | 9 | *Paracoccus*<br>*Sulfitobacter*<br>*Pseudoalteromonas*<br>*Bacillus*<br>*Clostridium*<br>*Desulfovibrio*<br>*Vibrio*<br>*Oscillatoria*<br>*Thalassospira* | ND | ND |
| Nelson et al.<br>(2011)<br>doi: 10.1038/ismej.2011.12 | Moorea, French Polynesia | Water associated from reef surrounds | 6 | *Acinetobacter*<br>*Pseudomonas*<br>*Pseudoalteromonas*<br>*Paracoccus*<br>*Bacillus*<br>*Pirellula* | ND | 0.0941 |
| Ainsworth et al.<br>(2015)<br>doi:10.1038/ismej.2015.39 | Great barrier reef and Hawaii | Coral associated - core microbiome | 13 | *Ralstonia*<br>*Bacteroides*<br>*Pseudomonas*<br>*Pseudoalteromonas*<br>*Corynebacterium*<br>*Limnobacter*<br>*Streptococcus*<br>*Halomonas*<br>*Staphylococcus*<br>*Acinetobacter*<br>*Stenotrophomonas*<br>*Vibrio*<br>*Psychrobacter* | 0.0029 | 0.1466 |

| | | | | | | |
|---|---|---|---|---|---|---|
| Ziegler et al. (2017) doi: 10.1038/ncomms14213 | Ofu Island, American Samoa | Coral associated | 5 | *Vibrio*<br>*Pseudomonas*<br>*Pseudoalteromonas*<br>*Brachybacterium*<br>*Arcobacter* | ND | ND |
| Hernandez-Agreda et al. (2016) doi: 10.1128/mBio.00560-16 | Great Barrier Reef and Coral Sea | Coral associated - core microbiome | 9 | *Pseudomonas*<br>*Mycobacterium*<br>*Vibrio*<br>*Alcanivorax*<br>*Staphylococcus*<br>*Ralstonia*<br>*Pseudoalteromonas*<br>*Acinetobacter*<br>*Corynebacterium* | 0.0024 | 0.1459 |
| **Mean match** | | | | | **2.92%** | **8.44%** |

ND = not detected.

**Table S3.** Relative abundance of the most abundant fungal ASV's at genus level to genera identified from the literature as components of coral associated microbiomes.

| **Publication** (see Supplementary Methods references list) | **Sample location** | **Microbiome type** | **Total number of matching genera** | **List of matching genera** | **Matching genera to continent air** | **Matching genera to ocean air** |
|---|---|---|---|---|---|---|
| Littman et al. (2011) | Great Barrier Reef | Coral associated | 1 | *Gibberella* | 0.0021 | ND |
| Morrison-Gardiner (2002) | Australian tropical waters | Sediment and marine organism related | 6 | *Alternaria*<br>*Cladosporium*<br>*Curvalaria*<br>*Epicoccum*<br>*Nigrospora*<br>*Periconia* | 0.1825 | 0.2577 |
| Sweet at al. (2013) | Caribbean | Coral associated and pathogens | 0 | | N/A | |
| Staley et al. (2017) | Caribbean | Coral associated and water associated proximate to reef | 1 | *Exophiala* | ND | 0.0042 |
| **Mean score** | | | | | **6.16%** | **8.73%** |

ND = not detected.
